# Gene Expression Associated with Individual Variability in Intrinsic Functional Connectivity

**DOI:** 10.1101/2021.06.01.446504

**Authors:** Liangfang Li, Yongbin Wei, Jinbo Zhang, Junji Ma, Yangyang Yi, Yue Gu, Liman Man Wai Li, Ying Lin, Zhengjia Dai

## Abstract

It has been revealed that intersubject variability (ISV) in intrinsic functional connectivity (FC) is associated with a wide variety of cognitive and behavioral performances. However, the underlying organizational principle of ISV in FC and its related gene transcriptional profiles remain unclear. Using resting-state fMRI data from the Human Connectome Project (299 participants) and microarray gene expression data from the Allen Human Brain Atlas, we conducted a transcription-neuroimaging association study to investigate the spatial configurations of ISV in intrinsic FC and their associations with gene transcriptional profiles. We found that the multimodal association cortices showed the greatest ISV in FC, while the unimodal cortices and subcortical areas showed the least ISV. Importantly, partial least squares regression analysis revealed that the transcriptional profiles of genes associated with human accelerated regions (HARs) could explain 31.29% of the variation in the spatial distribution of ISV in FC. The top-related genes in the transcriptional profiles were enriched for the development of the central nervous system, neurogenesis and the cellular components of synapse. Moreover, we observed that the effect of gene expression profile on the heterogeneous distribution of ISV in FC was significantly mediated by the cerebral blood flow configuration. These findings highlighted the spatial arrangement of ISV in FC and their coupling with variations in transcriptional profiles and cerebral blood flow supply.

## Introduction

Intersubject variability (ISV) in the functional connectivity (FC) of the human brain underlies individual differences in cognition and behavior (Finn and Todd Constable, 2016; Kelly et al., 2012; Smith et al., 2015). Recently, resting-state fMRI (R-fMRI) studies suggested that the ISV in intrinsic FC exhibited a sizeable regional variation for both adults (Li et al., 2019; Mueller et al., 2013) and neonates (Gao et al., 2014; Stoecklein et al., 2020), with higher ISV in multimodal association cortices than in unimodal cortices. These regions with high ISV in FC can not only predict individual differences in higher-order cognitive functions (e.g., inhibition and fluid intelligence) but also provide valuable information for individual identification (Finn et al., 2015; Horien et al., 2019; Liu et al., 2018). Moreover, previous works demonstrated that the overall pattern of ISV in FC detected in the neonatal brain was similar to the distribution observed in healthy adults (Gao et al., 2014; Stoecklein et al., 2020). These findings highlighted the effect of gene expression on the distribution of ISV in FC since the effect of the postnatal environment was limited during the neonatal period. In the present study, we aimed to investigate the mechanisms underlying the contributions of genetic factors to the distribution of ISV in FC.

Previous studies have begun to explore the genetic basis of human brain FC organization (Gao et al., 2014; Richiardi et al., 2015; Vértes et al., 2016; Wang et al., 2015). In the study of Gao et al. (2014), the genetic contribution to ISV in FC was estimated by comparing FC variability across monozygotic twins, dizygotic twins and unrelated singleton pairs. They found that an increased degree of genetic sharing (100% in monozygotic twins) was significantly associated with a decrease in FC variability, which indicated a strong genetic effect on ISV in FC. However, the previous study on the genetic contribution to ISV in FC mainly revealed the high heritability of FC, and which genes are associated with the distribution of ISV in FC remains unknown. Genes located in human accelerated regions (HARs) play a key role in neuron development, including cortical expansion, neurogenesis, and neuronal differentiation (Doan et al., 2016; Wei et al., 2019; Won et al., 2019). Neural changes during the neurogenesis processing are likely to be associated with the development of FC (Vogel et al., 2010), and FC has been found to be related to the genes associated with neuron development (Richiardi et al., 2015; Wang et al., 2015). Hence, we speculated that the gene expression profile of HAR genes would be a potential genetic underpinning of the distribution of ISV in FC. However, how the gene expression profile shapes the distribution of ISV in FC remains largely unknown.

The regions with high ISV in FC, which are primarily located in the prefrontal and parietal cortices, largely overlap with the regions with high cerebral blood flow (Liang et al., 2013). Resting-state cerebral blood flow (CBF) reflects the regional metabolism level and is a fundamental physiological property of the human brain (Satterthwaite et al., 2014). Moreover, resting-state CBF is also influenced by genetic factors that are involved in neurogenesis and neuron development (Goyal et al., 2014). In addition, a previous study has found that the genes that influence brain metabolism also regulate FC (Glahn et al., 2010). Hence, we speculated that the brain metabolism level might mediate the effect of gene expression on ISV in FC.

To uncover the mechanisms underlying the contributions of genetic factors to ISV in FC, we conducted a transcription-neuroimaging association study (Figure 1 for schematic of this study). Our first aim was to investigate the spatial configurations of ISV in FC based on high-resolution R-fMRI data from the Human Connectome Project (HCP; 299 participants) (Van Essen et al., 2013) and their relationship to a variety of cognitive abilities based on the meta-analysis method of the NeuroSynth database (Yarkoni et al., 2011). Based on previous studies (Gao et al., 2014; Mueller et al., 2013), we hypothesized that the high ISV in FC would be located in the association cortices, which tend to be responsible for higher-order functions. Our second aim was to investigate the relationship between gene expression profiles obtained from the Allen Human Brain Atlas (Hawrylycz et al., 2012) and intersubject variability in FC by directly examining the overlapping spatial variations of gene expression and ISV in FC. Since a previous study proposed that FC was influenced by genes related to neuron development (Wang et al., 2015), we hypothesized that the expression of HAR genes, which are crucial for brain neuron development, would significantly correlate with ISV in FC. Our third aim was to examine the potential mediation effect of resting-state CBF by conducting a mediation analysis to model the relationships among the gene expression profile, CBF, and the distribution of ISV in FC.

**Figure 1.**
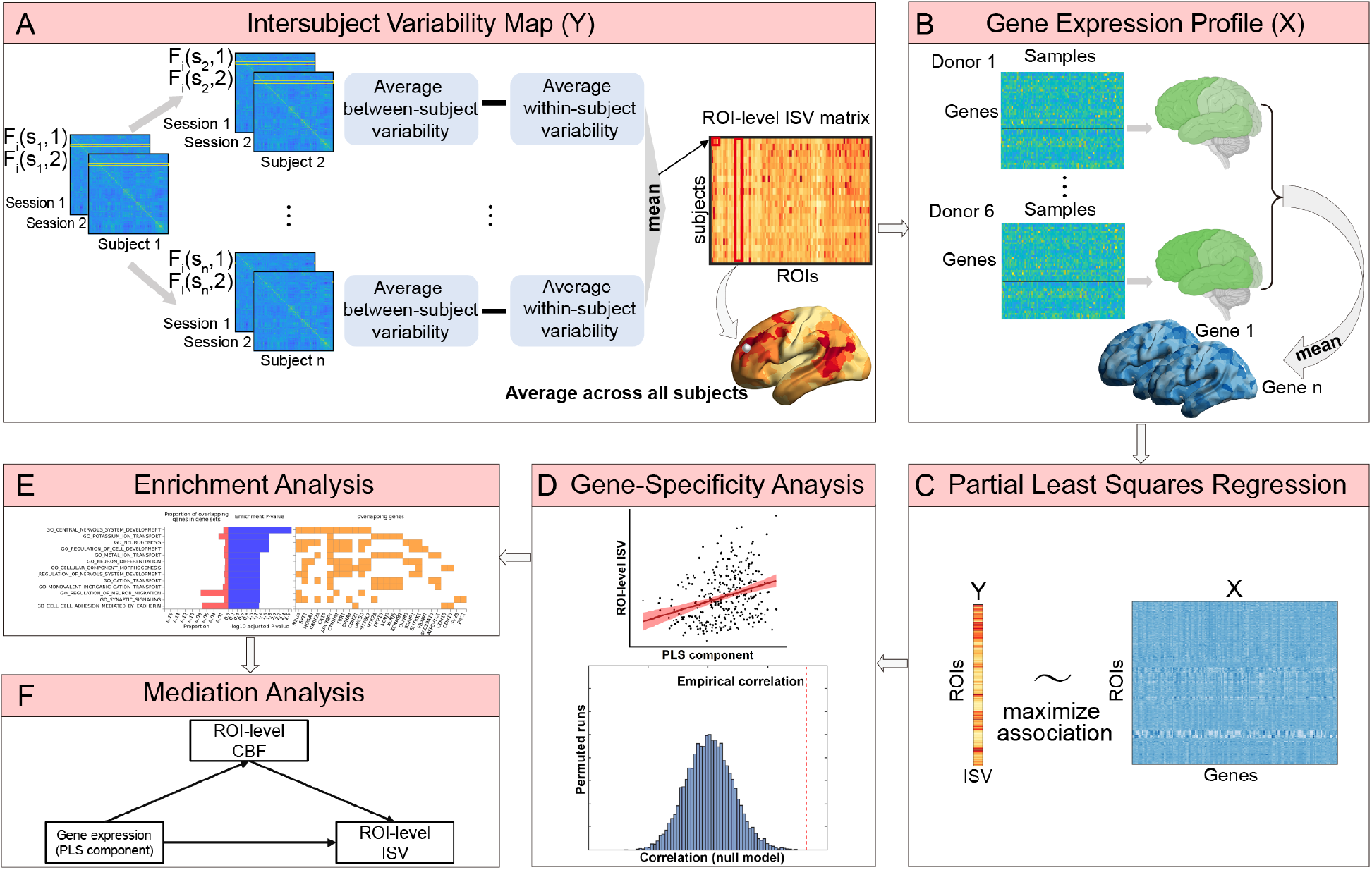
Schematic diagram of study design and methodology. (A) Using the repeat-measured R-fMRI data from HCP to calculate the ROI-level ISV in FC. (B) Using the microarray gene expression data across cortical regions from the Allen Human Brain Atlas to obtain the average gene expression profiles across six donors. (C) Using the partial least squares regression to investigate the association between the distribution of ISV in FC and gene expression profiles. (D) Gene-specificity analysis of ISV-related transcription-neuroimaging associations. (E) Gene enrichment analysis for top-related genes. (F) Mediation analysis for verifying the relationships between the genes, resting-state CBF and ISV in FC. ISV, intersubject variability; FC, functional connectivity; PLS, partial least squares; CBF, cerebral blood flow.

## Materials and Methods

### Participants

Data of 339 unrelated healthy adults from the released dataset of 900 participants were obtained from the Human Connectome Project (HCP; Van Essen et al., 2013). Since the HCP provides data from a large number of twins and non-twin siblings, we only selected unrelated participants, each with a unique family ID, to avoid confounding effects induced by shared genetic and environmental factors within a family structure. All participants were between 22 and 37 years old and had provided written informed content. The HCP project was approved by the Institutional Review Board of Washington University in St. Louis.

### R-fMRI data acquisition

All R-fMRI data were collected using a customized 32-channel Siemens 3T Connectome Skyra scanner. During scanning, the participants were asked to open their eyes, stare at the bright cross-hair on a black background, and relax. The R-fMRI data were collected in two sessions on two different days. Each session consisted of two run scans with a left-to-right (LR) and a right-to-left (RL) phase encoding direction, resulting in 4 resting-state run scans for each participant. Each R-fMRI run was acquired using a multiband gradient-echo-planar imaging sequence as follows: time repetition (TR) = 720 ms, time echo (TE) = 33.1 ms, flip angle = 52°, field of view = 208×180 mm^2^, matrix = 104×90, 72 slices, voxel size = 2×2×2 mm^3^, multiband factor = 8, and 1200 volumes (scanning time: 14.4 min). To eliminate the potential impact of different phase encoding directions on our findings, our analyses were restricted to the R-fMRI data with LR phase-encoding runs in two different sessions.

### R-fMRI Data preprocessing

The R-fMRI data were first preprocessed by the HCP minimal preprocessing procedure (Glasser et al., 2013), which included gradient distortion correction, head motion correction, image distortion correction, spatial transformation to the Montreal Neurological Institute (MNI) space, and intensity normalization. Forty participants were discarded due to excessive head motion in either session with the exclusion criteria of mean frame-wise head motion above 0.14 mm (Finn et al., 2015). Therefore, 299 participants (28.46±3.69 years old, 139 male/160 female) were used for the subsequent analyses. Further data preprocessing was performed using the Data Processing Assistant for Resting-State fMRI (DPARSF) (Yan and Zang, 2010; Yan et al., 2016) and Statistical Parametric Mapping software (SPM12; http://fil.ion.ucl.ac.uk/spm). These additional preprocessing steps included: (1) removing linear trend; (2) regressing out nuisance signals [including 24 head motion parameters (Friston et al., 1996), cerebrospinal fluid, white matter and global signals (Birn et al., 2006; Fox et al., 2009)]; and (3) performing temporal bandpass filtering (0.01–0.1 Hz). The residuals were used to construct FC matrix.

### FC matrix construction

To construct the whole-brain FC matrix, the parcellation atlas with 625 similar-sized regions was used to parcellate the brain gray matter (excluding cerebellum) into 625 regions of interest (ROIs), which preserved the automated anatomical labeling (AAL) landmarks (Dai et al., 2019; Tzourio-Mazoyer et al., 2002; Zalesky et al., 2010). The time series were then extracted from each ROI by averaging the time series of all voxels within that region. Finally, the Pearson correlation coefficients between the time courses of each possible pair of ROIs were calculated and normalized using Fisher’s *z*-transformation, resulting in a 625×625 FC matrix for each participant.

### ISV in FC

#### ROI-level ISV

To derive the spatial distribution map of ISV in FC across the whole brain, we calculated the ISV in FC pattern based on each ROI (Figure 1). The FC map of each ROI was denoted as a 624-dimensional real vector *F*_*i*_(*s, t*), where *i*∈{1, 2, …, 625}, *s*∈{1, 2, …, 299}, *t*∈{1, 2} indicated the respective indices of ROIs, participants, and scan sessions, and each element corresponded to the FC between ROI *i* and the remaining 624 ROIs. Given an ROI *i* and a participant *s*, the within-subject variability in ROI-level FC between two sessions was quantified as

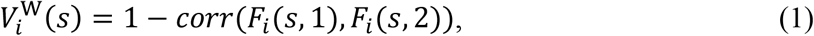

where *corr* was the function of the Pearson correlation. The average within-subject variability of two different participants *s*_1_ and *s*_2_ for ROI *i* was defined as

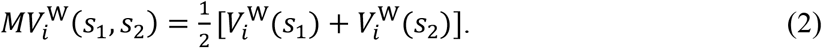

In addition to the within-subject variability, the between-subject variability between *s*_1_ and *s*_2_ for ROI *i* in each scan session was estimated as

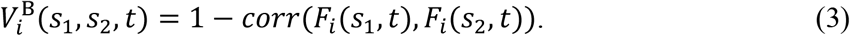

The average between-subject variability of two different participants across two sessions for ROI *i* was defined as

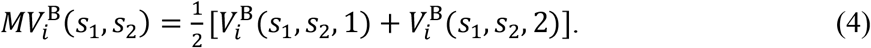

Based on the above definitions, we defined the intersubject variability (ISV) of ROI *i* between two different participants *s*_1_ and *s*_2_ by removing the average within-subject variability 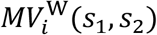 from the average between-subject variability 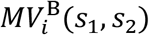, i.e.,

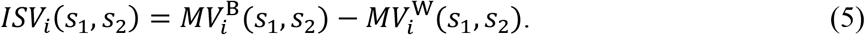

The intersubject variability *ISV*_*i*_ *(s)* of an ROI *i* regarding a single participant *s* was then calculated as the mean of the intersubject variabilities between *s* and the other 298 participants. By averaging the intersubject variabilities across all of the participants, we obtained the average ISV of ROI *i* as follows:

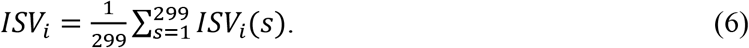

Finally, we repeated the above computation for all 625 ROIs, resulting in a 625×1 ISV map.

#### Within-module-level ISV

To quantify the ISV in FC at the modular level, the 625 ROIs were partitioned into seven functional modules (Yeo et al., 2011), including the visual (Vis), somatomotor (Mot), dorsal attention (dATN), ventral attention (vATN), limbic (LMB), frontoparietal (FPN), and default mode network (DMN). For each module, we first extracted the FC values between all pairs of ROIs within the module to produce the within-module FC maps. Then, the ISV of the within-module FC patterns was calculated in a similar way as that of the above ROI-level ISV, except that the FC map of each ROI was replaced by the within-module FC map (for more details, please refer to the Supplemental Information). As such, we obtained a 299×7 within-module-level ISV matrix.

To compare the mean within-module-level ISV difference between different modules, a nonparametric permutation test (N = 10000) was performed. The false discovery rate (FDR) correction was adopted for multiple comparisons with a significance threshold of *p* < 0.05.

### Behavioral data

To investigate how the regional ISV in FC could be associated with cognitive topic terms, a NeuroSynth meta-analysis was implemented using the NeuroSynth meta-analytic database (http://www.neurosynth.org) (Yarkoni et al., 2011). Specifically, the original ISV map was split into five-percentile increments and then binarized to obtain 20 binary maps. Each of the 20 binary maps was used as an input to the meta-analysis, and the outputs of the analysis were *z*-statistics associated with the 23 topic terms representing a wide range of behaviors (Margulies et al., 2016; Preti and Van De Ville, 2019).

### Gene expression

#### AHBA dataset

To characterize the genetic underpinnings of ISV in FC, the gene symbol list of 415 brain-expressed HAR genes (further referred to as HAR-BRAIN genes) was first obtained from Wei et al. (2019), and the gene expression profiles of these HAR-BRAIN genes were extracted from the complete microarray gene expression data from the Allen Human Brain Atlas dataset (AHBA) (http://human.brain-map.org/static/download) (Hawrylycz et al., 2012). AHBA consists of expression profiles of 20700 genes for 3702 spatially distinct tissue samples collected from the post-mortem brains of six human donors (all without a history of neuropsychiatric or neuropathological disorders) (for details, see Table S1). Tissue samples of the left hemisphere were included in this study, as data from all six donors were available for the left hemisphere, and only two donors were available for the right hemisphere.

The samples in AHBA were first mapped into ROIs by computing the minimal Euclidean distance between each sample and all gray matter voxels located in the left hemisphere of the AAL-625 atlas (for more details, please refer to the Supplemental Information). The gene expression data were computed for each ROI by averaging the expression data of the samples mapped to that particular ROI. Gene expression data of each gene were normalized to *z*-scores across all ROIs in each donor’s dataset. Normalized gene expression data were then averaged across six donors to obtain a group-level gene expression matrix. Gene expression data of 415 HAR-BRAIN genes were extracted from the complete group-level gene expression data matrix. To investigate whether the HAR-BRAIN genes were more highly expressed particularly in the modules with high within-module-level ISV (e.g., FPN, DMN, dATN), a nonparametric permutation test (N = 10000) with FDR for multiple comparisons correction was performed to compare the mean gene expression difference between modules.

### Relationship among ISV in FC, HAR-BRAIN gene expression, and CBF

#### Correlation between ISV map and HAR-BRAIN gene expression profile

To verify our hypothesis that the HAR-BRAIN gene expression may be a genetic root of ISV in FC, we used partial least squares (PLS) regression to identify the expression patterns of HAR-BRAIN genes that were significantly correlated with ISV in FC. PLS regression is a multivariate analysis aiming to identify the components, which were linear combinations of the weighted gene expression scores (predictor variables), that were the most predictive to ISV in FC (response variables). PLS regression has been widely used for neuroimaging and transcriptional data analyses (Liu et al., 2020; Morgan et al., 2019; Seidlitz et al., 2018; Vértes et al., 2016; Whitaker et al., 2016).

To examine whether the PLS components can significantly explain the variation in ISV of FC, we adopted a permutation test with the spatial autocorrelations corrected by generative modeling (Burt et al., 2020) to examine whether the real R^2^ of the component that explained more than 10% of the total variance (Liu et al., 2020) was significantly larger than that achieved by chance. We also used the permutation test with spatial autocorrelation corrected to examine the significance of the empirical spatial correlation between PLS components and the ROI-level ISV map. Moreover, to determine whether the HAR-BRAIN genes were more specifically associated with the ISV map than the other genes, we randomly selected 415 genes from the pool of 2564 BRAIN genes (excluding HAR-BRAIN genes) (Wei et al., 2019) to repeat the PLS regression 10000 times. The Spearman correlation between the PLS components and the ISV map obtained from 10000 repetitions constituted a null model. Furthermore, we used the bootstrapping method to estimate the error in the estimation of the weight of each gene and divided the weight of each gene by the estimated error to generate the corrected weight. We ranked the genes according to their corrected weights, which represented their contributions to the PLS component.

To identify the possible biological functions of ISV-related genes, we performed gene-enrichment analysis for the top-ranked related genes with positive weight (top 10%) in the first few significant components by using the hypergeometric test implemented in the GENE2FUNC function of FUMA (https://fuma.ctglab.nl) (Watanabe et al., 2017). This test examined whether the top-ranked related genes were enriched for the predefined Gene Ontology (GO) gene sets that have specific functional interpretations in three functional categories, including biological process, cellular component, and molecular function. For each of the predefined GO gene sets, a *p*-value was calculated based on the number of genes present in both the predefined set and the top-ranked related gene set. The resulting *p*-values were corrected for multiple testing through FDR with *p* < 0.05. The GO terms with an FDR *p*-value below 0.05 were reported.

#### Correlation between ISV map and the CBF map

To explore whether the spatial distribution of ISV in FC can reflect the CBF configuration, we compared the spatial distributions of the ISV map and those of the CBF map in the resting state, which shows the brain regions’ metabolic costs (Satterthwaite et al., 2014). We calculated the spatial correlation between the ROI-level ISV map and CBF map and performed spatial autocorrelation-correcting permutation test to examine the significance of the empirical spatial correlation.

#### Mediation analysis

To test the hypothesis that the CBF distribution mediated the contribution of the gene expression profile to the ISV map, a bootstrapped mediation analysis was performed using the simple mediation model from the PROCESS macro in SPSS (Hayes, 2017). The mediation analysis was conducted with 5000 bootstrap samples to generate bias-corrected confidence intervals (CI). The indirect effect was considered significant when the bootstrapped 95% CI did not include zero (Hayes, 2017).

## Results

### Spatial distribution of ISV in FC across brain regions and intrinsic modules

We observed that the spatial distribution of ISV in FC [measured by Eq. (6)] was regionally heterogeneous (Figure 2A). The association cortices, including the prefrontal (dorsolateral superior frontal gyrus, middle frontal gyrus, inferior frontal gyrus), temporal (middle temporal gyrus, superior temporal gyrus), and parietal lobe (supramarginal gyrus), showed high ISV. Meanwhile, the unimodal cortices, including the primary visual (cuneus, lingual gyrus, superior occipital gyrus), sensorimotor (postcentral gyrus, precentral gyrus), and subcortical areas (pallidum, caudate nucleus, thalamus, amygdala) showed low ISV. This pattern was compatible with previous observations of individual variability in FC (Mueller et al., 2013).

**Figure 2.**
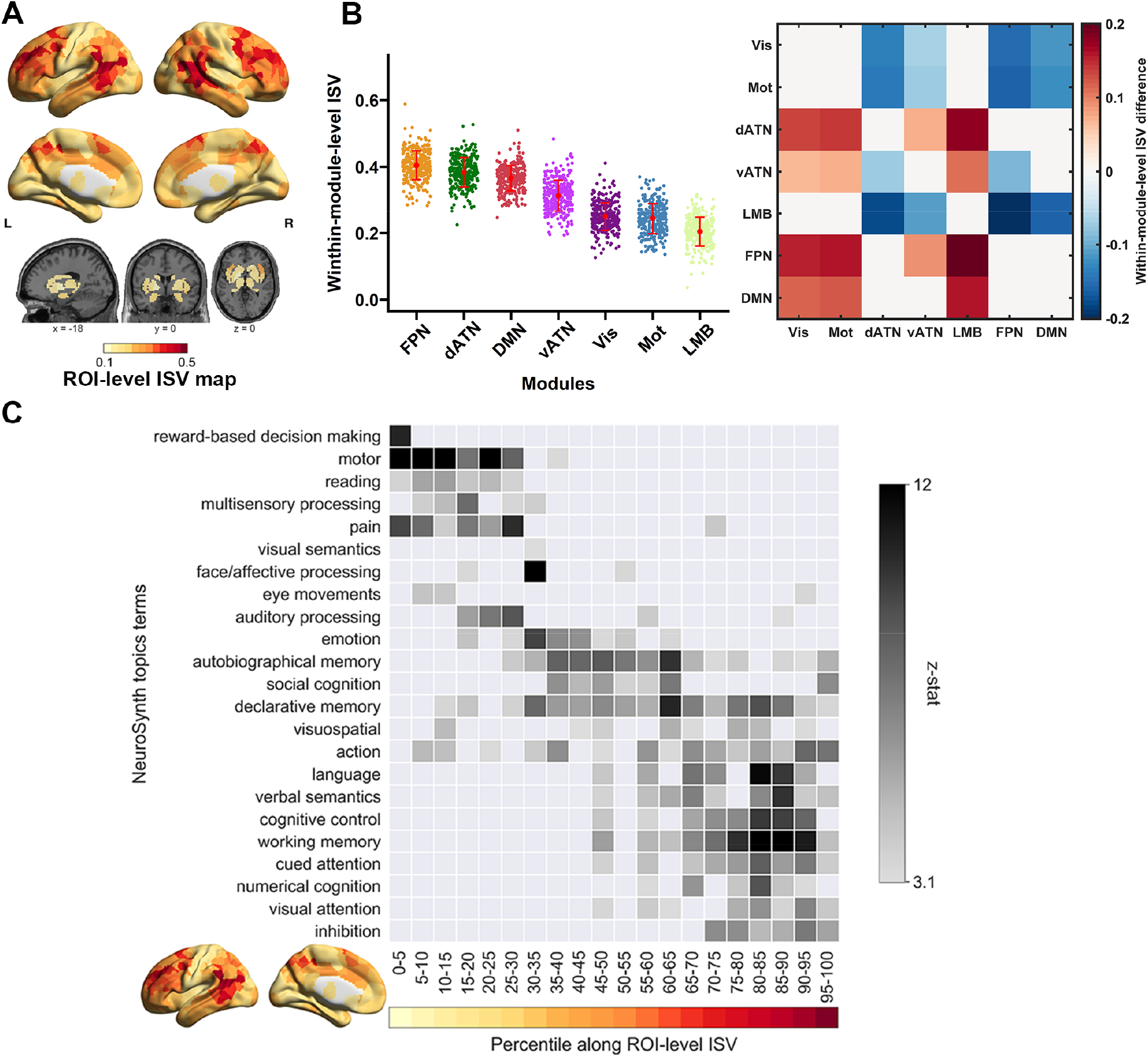
Spatial distribution map of ISV in FC and results of NeuroSynth meta-analysis. (A) Spatial distribution of ROI-level ISV. The subcortical regions displayed in Figure 2 and 3 included insula, caudate, putamen, pallidum, thalamus, hippocampus, and amygdala. (B) Comparison of within-module-level ISV. The error bar indicates the standard deviation (SD) of the within-module-level ISV across all the participants under each module. The matrix on the right shows the within-module-level ISV differences between the modules (row-column); within-module-level ISV differences that are not significant in the permutation test (*p* > 0.05) are set to zeros. (C) NeuroSynth meta-analysis of regions of interest along the ISV using 23 topic terms. Terms are ordered by the weighted mean of their location along the ISV. ISV, intersubject variability; FC, functional connectivity.

For within-module-level ISV, visual inspection indicated that higher-order cognitive modules, including FPN and dATN, had the highest variability, whereas LMB and the primary modules, including the Vis and Mot, had the lowest variability (Figure 2B) (for details, see Table S2). The permutation test revealed that the ISV of the four higher-order cognitive modules (FPN, dATN, DMN, vATN) was significantly higher than that of LMB and the primary modules (Vis, Mot) (*p*s < 0.035, 10000 permutations, FDR correction). In addition, the ISV of vATN was significantly lower than that of FPN and dATN (*p*s < 0.024, 10000 permutations, FDR correction) (for details, see Table S3).

### Spatial distribution of ISV in FC reflected a cognitive spectrum from primary to higher-order functions

By overlapping the 20 binary ISV maps with the 23 cognitive term maps available from the NeuroSynth database, we found that low-ISV regions were more related to primary functions, such as “motor”, “multisensory processing”, and “auditory processing”, and high-ISV regions were related to higher-order functions, such as “language”, “cognitive control”, “working memory”, “numerical cognition” and “inhibition” (Figure 2C). The behavior of the topics arranged from bottom to top corresponded to the function of the modules in Figure 2B that was sorted according to the within-module-level ISV.

### Expression profile of HAR-BRAIN genes across brain regions and intrinsic modules

To investigate how the spatial distribution of ISV in FC was related to the expression of 415 HAR-BRAIN genes, we mapped the samples in the AHBA dataset to the ROIs, and estimated the average expression of 415 HAR-BRAIN genes for each ROI. HAR-BRAIN genes showed high-level expression in regions of the frontal cortices (medial superior frontal gyrus, orbital superior frontal gyrus, dorsolateral superior frontal gyrus), temporal pole (superior temporal gyrus, middle temporal gyrus), and anterior cingulate and paracingulate gyri. Meanwhile, the visual cortices (fusiform gyrus, lingual gyrus) and subcortical regions (thalamus, pallidum, parahippocampal gyrus, caudate nucleus, putamen) displayed low-level gene expression (Figure 3A). Hence, there was an overall tendency that the average expression of HAR-BRAIN genes increased from subcortical regions and primary areas to the association cortices.

**Figure 3.**
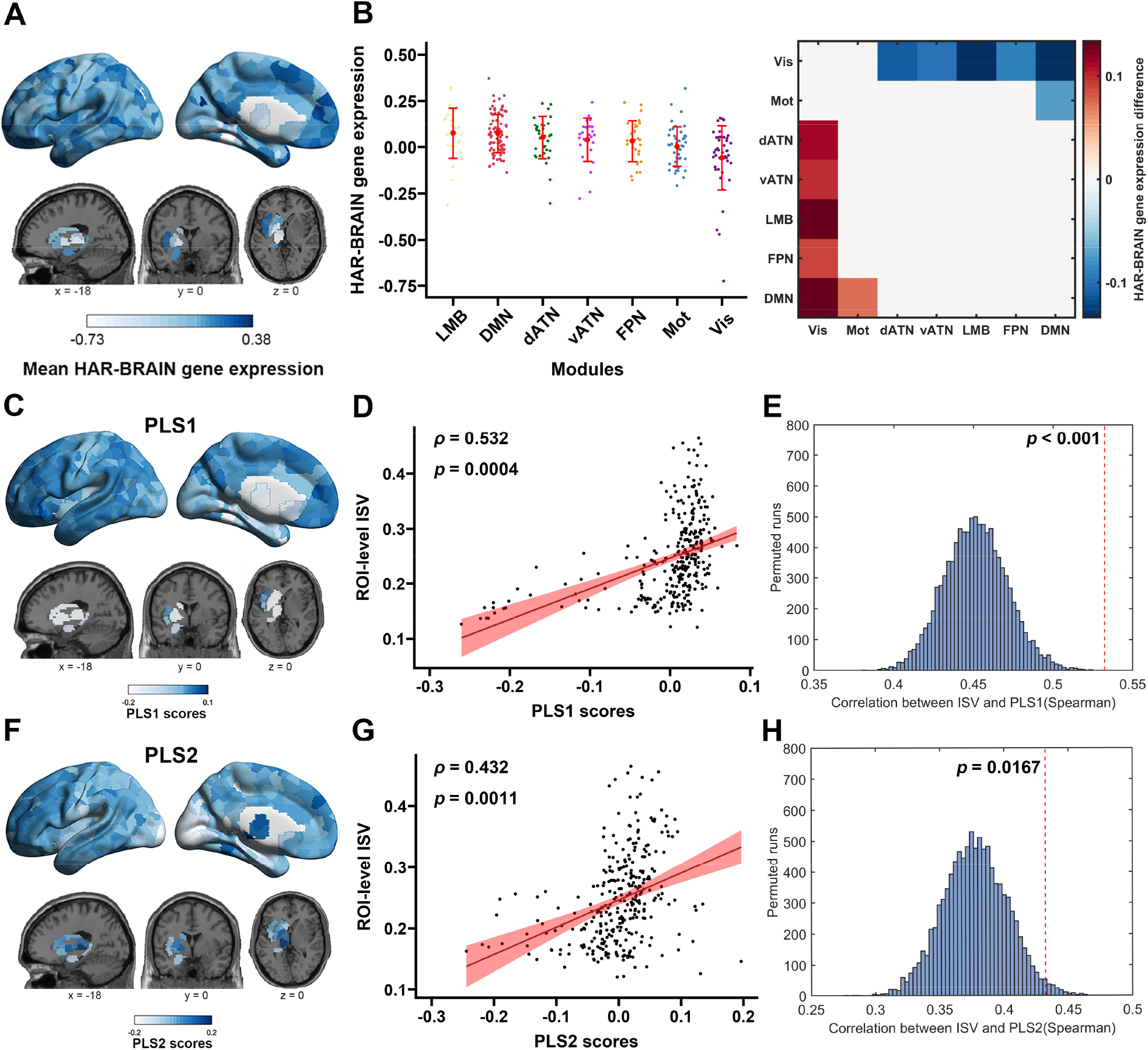
Association between gene expression profiles and ISV in FC. (A) Average expression profile of HAR-BRAIN genes. (B) HAR-BRAIN gene expression within each module ranked in the descending order of the mean gene expression. The error bar indicates the SD of the average gene expression levels of ROIs within the same module. The matrix on the right shows the gene expression differences between the modules (row-column); gene expression differences that are not significant in the permutation test (*p* > 0.05) are set to zeros. (C) PLS1 identifies a gene expression profile with overexpression mainly in the prefrontal, parietal and lateral temporal cortices. (D) Spearman correlation between PLS1 and ROI-level ISV in FC. The red shadow indicates the 95% CI. (E) Gene-specificity analysis of the association between PLS1 and ROI-level ISV in FC. The red dotted line represents the empirical correlation value. (F) PLS2 identifies a gene expression profile with overexpression dominantly in the subcortical, temporal and prefrontal cortices. (G) Spearman correlation between PLS2 and ROI-level ISV in FC. (H) Gene-specificity analysis of the association between PLS2 and ROI-level ISV in FC. ISV, intersubject variability; FC, functional connectivity; PLS1, the first partial least squares regression component; PLS2, the second partial least squares regression component.

The permutation test indicated that the HAR-BRAIN genes showed significantly higher expression levels in LMB, DMN, dATN, vATN, and FPN than in Vis (*p*s < 0.024, 10000 permutations, FDR corrected), but no significant difference was detected among these five modules (for details, see Table S5). In addition, the gene expression in the DMN was significantly higher than that in the Mot (*p* = 0.024, 10000 permutations, FDR corrected) (Figure 3B).

### Gene expression profile was associated with the spatial distribution of ISV in FC

Using PLS regression, we found that two significant components explained 31.29% of the variance in ISV (*p* < 0.001, permutation tests with spatial autocorrelation corrected, 10000 permutations) (Figure S1 and S2). Specifically, the first partial least squares component (PLS1) represented a significantly positive association between ISV in FC and a gene expression profile characterized by high expression mainly in the prefrontal, parietal, and lateral temporal areas (Spearman’s ρ = 0.532, *p* = 0.0004, permutation tests with spatial autocorrelation corrected, 10000 permutations) (Figure 3C, 3D). The second independent partial least squares component (PLS2) also represented a significantly positive association between ISV in FC and a gene expression profile displaying high expression predominantly in the subcortical, temporal and prefrontal areas (Spearman’s ρ = 0.432, *p* = 0.0011, permutation tests with spatial autocorrelation corrected, 10000 permutations) (Figure 3F, 3G). The results of the gene-specificity analysis showed that the Spearman correlation between PLS1, PLS2, and ISV in FC was significantly higher than that of the null model (PLS1: *p* < 0.001; PLS2: *p* = 0.0167), which was based on equally sized random gene sets taken from BRAIN genes (Figure 3E, 3H). These findings suggested that the HAR-BRAIN genes played a specific role in shaping the spatial distribution of ISV in FC.

Moreover, the transcriptional profile of PLS1 was significantly enriched in genes related to the biological processes involved in central nervous system development, regulation of cell development, neurogenesis and potassium ion transport (all *p* < 0.05) (Figure 4A), and the transcriptional profile of PLS1 was significantly enriched in genes associated with the cellular components of the synapse (presynapse and postsynapse), synaptic membrane, membrane protein complex, glutamatergic synapse and cation channel complex (all *p* < 0.05) (Figure 4B). Meanwhile, the transcriptional profile of PLS2 was significantly enriched with genes related to the cellular component of the synaptic cleft (all *p* < 0.05) (Figure 4C).

**Figure 4.**
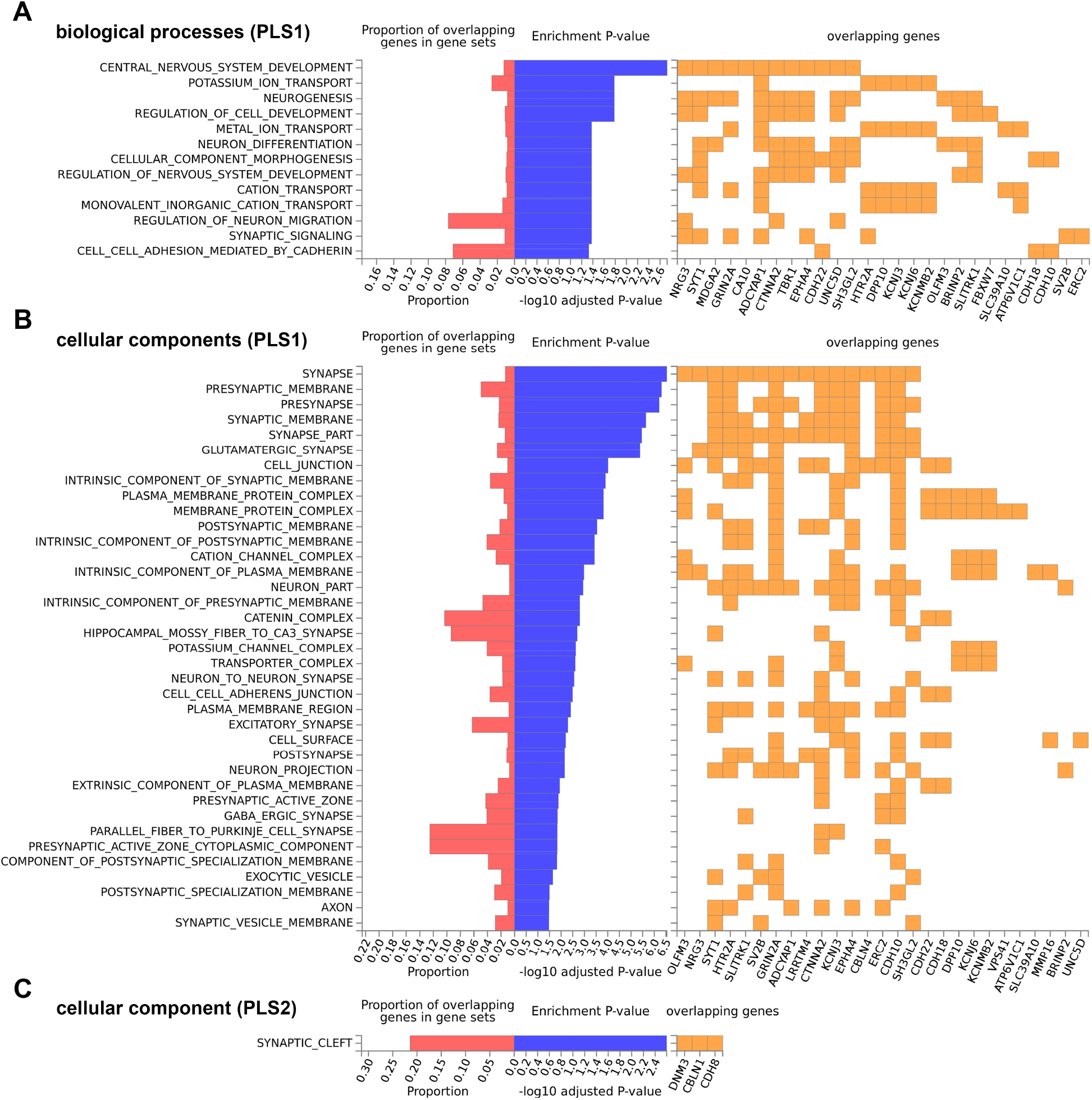
FUMA results of the overrepresented gene sets retrieved from the functional enrichment analysis of the top-ranked related genes in PLS1 and PLS2. The blue bar indicates the enrichment *p*-value (−log10 adjusted) after FDR correction. The red bar indicates the proportion of overlapping inputted genes according to the size (number of genes) of each gene set. The orange square indicates the inputted genes that are included in the enriched gene sets. (A) Significantly enriched gene sets from the Gene Ontology—biological process category for PLS1. (B) Significantly enriched gene sets from the Gene Ontology—cellular component category for PLS1. (C) Significantly enriched gene sets from the Gene Ontology—cellular component category for PLS2. PLS1, the first partial least squares regression component; PLS2, the second partial least squares regression component.

### CBF mediated the association between the gene expression profile and ISV in FC

As reported by Satterthwaite et al. (2014), brain CBF varied regionally in the resting state, with high CBF primarily distributed in the bilateral dorsolateral prefrontal cortex, superior and medial frontal cortex, posterior cingulate cortex, lateral temporal cortex and inferior parietal lobes (Figure 5A). A significant positive correlation was shown between the CBF and ISV in FC (Spearman’s ρ = 0.336, *p* = 0.0034, permutation tests with spatial autocorrelation corrected, 10000 permutations) (Figure 5B). Additionally, we found that higher gene expression of PLS1 was associated with higher CBF (path a: β = 0.313, *p* < 0.001). After controlling the influence of PLS1, higher CBF was related to higher ISV (path b: β = 0.201, *p* < 0.001). After considering the effect of CBF, the effect of PLS1 on ISV was weakened (path c’: β = 0.378, *p* < 0.001, from path c: β = 0.441, *p* < 0.001). Furthermore, mediation analysis revealed that CBF was a significant mediator (indirect effect = 0.063, 95% CI = [0.031, 0.104]), partially explaining the positive association between PLS1 and ISV (Figure 5C, top), while CBF did not mediate the positive association between PLS2 and ISV (indirect effect = 0.019, 95% CI = [−0.008, 0.050]) (Figure 5C, bottom).

**Figure 5.**
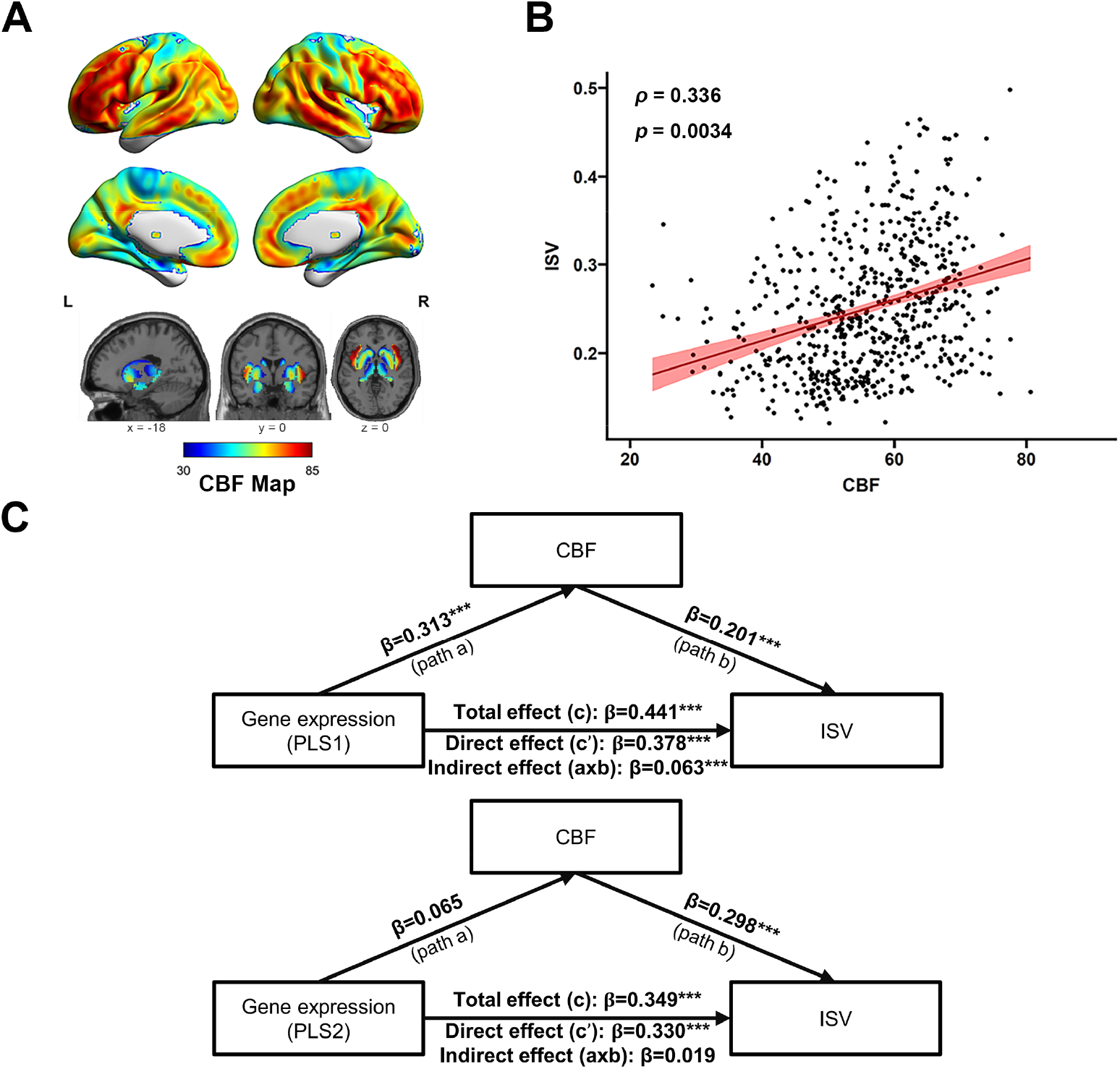
The CBF map and mediation analysis. (A) CBF map extracted from Satterthwaite et al. (2014). (B) Spearman correlation between ROI-level CBF and ROI-level ISV. The red shadow indicates the 95% CI. (C) The relationship between gene expression profiles of PLS1 (top) and ISV in FC was mediated by CBF. Standardized regression coefficients were reported. CBF, cerebral blood flow; ISV, intersubject variability; PLS1, the first partial least squares regression component; PLS2, the second partial least squares regression component. ^***^*p* < 0.001.

## Discussion

Using R-fMRI, gene expression, and CBF data, we showed that the changes in the human genome during evolution played an important role in shaping the distribution of ISV in FC. First, we found that ISV in FC distributed heterogeneously across the whole brain, showing greater ISV in multimodal association cortices whilst less ISV in unimodal cortices and subcortical areas. Additionally, we found that the distribution of ISV in FC reflected a cognitive spectrum from primary to higher-order functions using a NeuroSynth meta-analysis. Second, we demonstrated that the distribution of ISV in FC was correlated with the transcriptional profiles of HAR-BRAIN genes, and the most correlated genes were related to the development of synapses and the central nervous system, neurogenesis, and neuron differentiation. Finally, we revealed that the effect of the gene expression profile on the heterogeneous distribution of ISV in FC was significantly mediated by the CBF configuration. Together, these findings may enhance our understanding of the molecular and neural mechanisms associated with the spatial arrangement of ISV in FC.

Several limitations of the current study should be considered. First, the gene expression data from the AHBA were sampled from six donors who had different ethnicities and sexes. This limited sample might have created a bias in capturing the variance in gene expression across individuals. However, AHBA is currently the only database that can provide whole-brain mRNA gene expression data. Second, the currently reported variability map was constructed based on the imaging data of living participants, while the gene expression data were derived from post-mortem brains, and the CBF map was constructed based on another dataset. Therefore, the results may be influenced by differences among the datasets, making it difficult to capture the relationship among the gene expression profile, CBF, and ISV in FC at the individual level. Future studies could implement individual-level genome-wide analysis and metabolic data to help further understand the genetic and physiological basis underlying ISV in FC.

